# Dispersal of southern elephant seals (*Mirounga leonina*) from Davis Base, Antarctica: Combining population genetics and tracking data

**DOI:** 10.1101/2021.11.26.470169

**Authors:** Michelle Chua, Simon Y. W. Ho, Clive R. McMahon, Ian Jonsen, Mark de Bruyn

## Abstract

Marine animals such as the southern elephant seal (*Mirounga leonina*) rely on a productive marine environment and are vulnerable to oceanic changes that can affect their reproduction and survival rates. Davis Base, Antarctica, acts as a moulting site for southern elephant seals that forage in Prydz Bay, but the genetic diversity and natal source populations of these seals has not been characterized. Determining the genetic diversity of moulting populations like this one provides essential information on seal dispersal, inter-population mixing, and foraging behaviours. In this study, we combined genetic and animal tracking data on these moulting seals to identify levels of genetic diversity, natal source population, and movement behaviours during foraging and haul-out periods. Using mitochondrial sequence data, we identified two major breeding lineages of seals at Davis Base. We found that the majority of the seals originated from breeding stocks within the South Atlantic Ocean and South Indian Ocean. One seal was grouped with the Macquarie Island breeding stock (South Pacific Ocean). The Macquarie Island population, unlike the other two stocks, is decreasing in size. Tracking data revealed long-distance foraging activity of the Macquarie Island seal around Crozet Islands. We speculate that changes to the Antarctic marine environment have resulted in a shift in foraging and dispersal strategies, which subsequently affects seal population growth rates. These findings have implications for conservation management plans aimed at improving the population status of the southern elephant seal.

## Introduction

Warmer oceans result in loss of sea ice, which is likely to have large impacts on reproduction and survival rates in species that depend on sea ice for foraging (Bryndum-Buchholz et al. 2019). In other species, loss of sea ice has reduced barriers to dispersal, which has resulted in geographical isolation of some marine animal populations (Laidre et al. 2018). As global temperatures continue to increase, changes to habitat structure in ice-locked regions are likely to cause shifts in marine animal movements, effective dispersal, foraging behaviours, and population numbers (Hindell et al. 2017; Hindell et al. 2020). These changes can lead to a restructuring or even loss of genetic diversity between populations (Laidre et al. 2018; Siegert et al. 2019).

Antarctic sea ice increased from the late 1970s to a peak in 2014, but large declines in 2017 and 2018 were possibly caused by the El Niño Southern Oscillation (Parkinson 2019). Polar marine animals, such as pinnipeds, occupy the upper trophic levels in the Antarctic region and therefore integrate large environmental signals (Bestley et al. 2020). Sea ice is crucial to the survival of these predators, especially those that require sea ice for breeding, foraging, and moulting (Bestley et al. 2020). A recent study using a climate model projected that the colonies of many Antarctic species, including emperor penguins, would become quasi-extinct by 2100 because of the reduced availability of sea ice and foraging habitat (Jenouvrier et al. 2021). Similarly, changes in the extent of sea ice have altered the foraging behaviour and survival rates of a number of pinnipeds, such as the southern elephant seal (*Mirounga leonina*), which has ultimately affected the population dynamics of these species (McMahon and Burton 2005; Bestley et al. 2020; Bester 2021).

Southern elephant seals (SES) have a circumpolar distribution and haul out twice a year on sub-Antarctic islands to breed and to moult (Le Boeuf and Law 1994). The four main breeding stocks are South Georgia, Kerguelen Islands and Heard Island, Macquarie Island, and Península Valdés in Argentina (McMahon et al. 2017), with smaller breeding colonies on other sub-Antarctic islands (Slade et al. 1998; Hoelzel et al. 2001) (Fig. 1). Population decreases of SES were first documented in the mid-1980s (McMahon et al. 2005a). Population estimates between the 1980s and early 2000s demonstrated stable and increasing population sizes across three of the main breeding stocks and most of the smaller breeding stocks (McMahon et al. 2005a). However, the Macquarie Island breeding stock has continued to decrease and has been listed as vulnerable due to dramatic decreases in population size (McMahon et al. 2005a; van den Hoff and Burton 2007). Initial decreases were likely to have been driven by hunting for the seal’s oil-rich blubber in the early 19th Century (Hindell and Burton 1988: van den Hoff and Burton 2007), but recent population declines were most likely caused by changes in food supply (McMahon et al. 2005a).

**Fig. 1.**
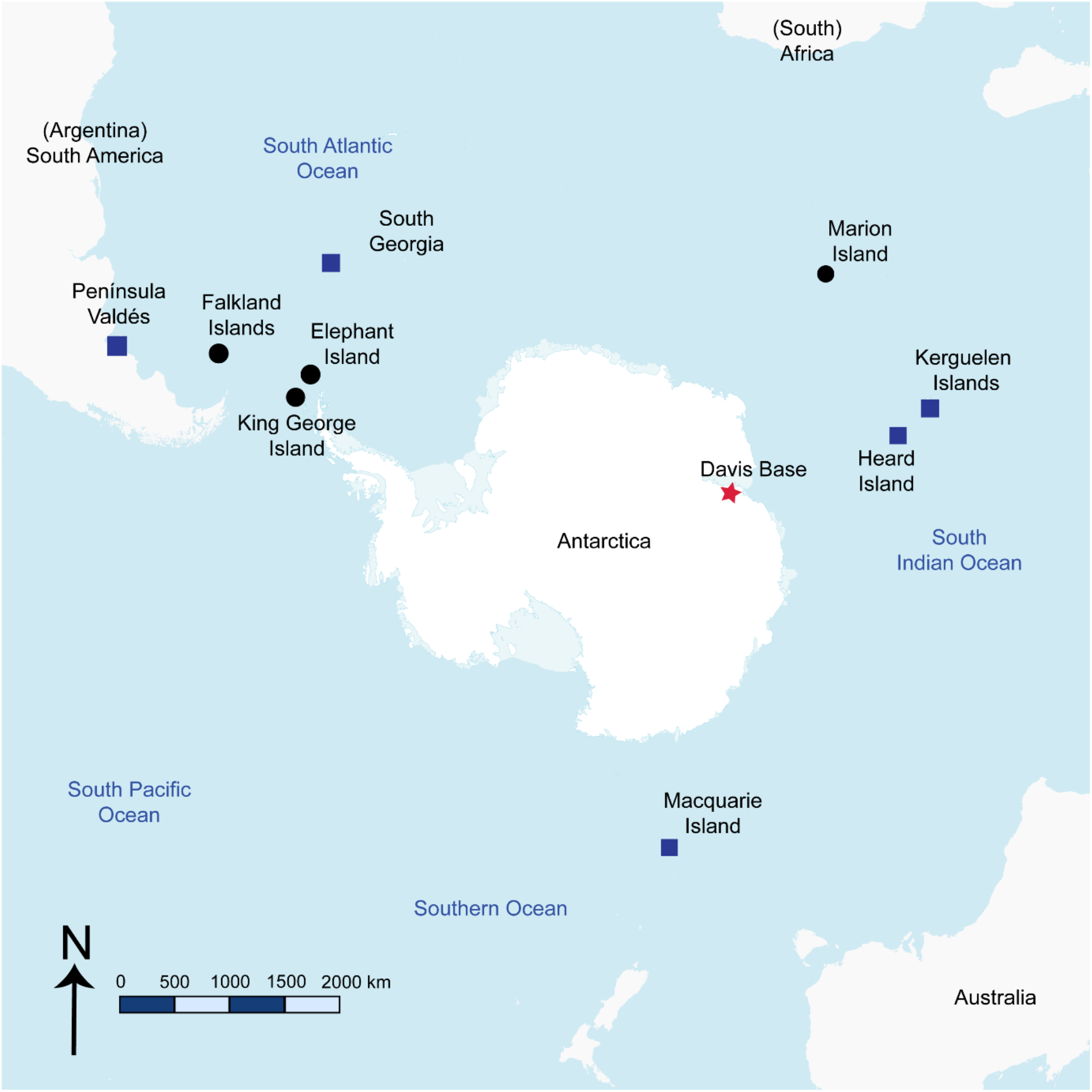
Circumpolar distribution of southern elephant seals around Antarctica. Blue squares represent the four main breeding stocks (South Georgia, Kerguelen Islands and Heard Island, Macquarie Island, and Península Valdés in Argentina). Black circles represent smaller breeding colonies (Marion Island, King George Island, Elephant Island, and Falkland Islands). The red star represents the sample collection site (Davis Base, Antarctica) for this study. Outline of continents and island locations are from Free Vector Maps.

During the 2.5-month period between the breeding and moulting seasons, and the eight-month period between moulting and breeding, SES spend the majority of their time foraging at sea. The timing and location of the seals’ foraging movements depend on energy requirements and the availability of food sources (Goedegebuure et al. 2018; de Kock et al. 2021). Polar research stations such as Casey Station (66° 16′ 57″ S, 110° 31′ 36″ E) and Davis Base (68° 34′ 36″ S, 77° 58′ 03″ E) act as moulting sites for some seal colonies (Rodríguez et al. 2017). Davis Base is an ice-free area that is situated near Vestfold Hills, covering roughly 400 km^2^ (Australian Antarctic Program 2020). Previous studies indicated that the male seals at Davis Base originated from the Kerguelen Islands, Heard Island, and some of the smaller populations in the Kerguelen stock at Marion and Crozet islands (Tierney 1977; Bester 1988). The seals’ annual visits might be due to the abundance of food along Prydz Bay during the austral summer (Bester 1988). The underlying genetic make-up of this and indeed other moulting aggregations and the possibility of sex-biased gene flow can potentially be identified through genetic analysis. The results of such an analysis would allow comparison with traditional capture-mark-recapture studies that formed the basis of much previous work on population composition at Davis Station.

The genetic structure of the breeding colonies of SES have been investigated previously (Hoelzel et al. 1993; Slade et al. 1998; Fabiani 2002; Chauke 2008; Bogdanowicz et al. 2013). However, there remains a gap in our knowledge regarding the natal origins of seals at moulting sites such as Davis Base, where seals potentially from multiple breeding sites aggregate (Bester 1988). Since SES are polygynous and display philopatry, breeding colonies have high female-to-male ratios (one male beachmaster to a female harem of up to 100 seals) (Bogadanowicz et al. 2013). Therefore, the reproductive success and dispersal of a single male can affect disproportionately the genetic structure of an entire breeding colony. These genetic effects on population composition can be investigated using molecular markers such as mitochondrial DNA (mtDNA), which is maternally inherited. Sequences of the mitochondrial control region effectively delineate the major breeding stocks of SES (Hoelzel et al. 1993; Slade et al. 1998; Bogadanowicz et al. 2013).

In this study, we aim to resolve the natal origins of SES that haul out at the moulting site at Davis Base, Antarctica. We analyse mtDNA from blood samples and tracking data from these seals to determine their genetic diversity and dispersal patterns, and draw comparisons with previous studies of the genetic structure of surrounding breeding colonies. Our study shows that genetic and physical tracking data provide complementary information on the natal affiliations of the seals and their resource usage around the Kerguelen Plateau.

## Materials and methods

### Blood sample collection and animal tracking tags

We captured, collected blood samples from, and attached animal tracking devices to 12 male SES (10 subadults and 2 juveniles) at Davis Base in 2016 and 2017. Animal tracking data were collected over one year of the seals’ lifecycle, from the start of their trip (moulting at Davis Base) to returning to moult the following year. After collection, blood from each sampled seal was spotted onto Whatman FTA cards (Stowell et al. 2018) and stored at −20°C until required for DNA extraction. The 12 blood samples were then sent to the University of Sydney for the purpose of this study.

### DNA library assembly

We prepared seal blood samples using the standard sterile technique, which involved taking a sterilized (Bunsen flame) standard office one-hole punch from the centre of the dried blood sample (Stowell et al. 2018). We extracted and purified the sampled DNA using the Qiagen DNeasy Blood & Tissue Kit following the Spin-Column protocol for blood with non-nucleated erythrocytes (Supplementary Information S1). To amplify mtDNA (Supplementary Information S2), we used the primers ancF (5′-GCTGACATTCTACTTAAACT-3′) and mdbR (5′-CAGTATAGAAACCCCCACATGA-3′) (de Bruyn et al. 2009). We ran PCRs for approximately 3 h using the following cycle protocol: 10 min at 95°C followed by 35 cycles of 30 s at 52°C, 30 s at 72°C, 1 min at 72°C, followed by an additional extension step for 10 min, and then 4°C for cooldown. After successful PCR, we cleaned up the PCR product following the ExoSAP-IT Express PCR Product Cleanup kit and standard protocol. The cleaned PCR products were stored at 4°C prior to DNA sequencing by Macrogen (Seoul, South Korea).

We collected a total of 201 additional sequences of the mitochondrial control region from GenBank (Hoelzel et al. 1993; Slade et al. 1998; Fabiani 2002; Chauke 2008; Curtis et al. 2009; Bogdanowicz et al. 2013; Zappes et al. 2017). These comprised 191 sequences from southern elephant seals and 10 sequences from Weddell seals (*Leptonychotes weddellii*),which we included as an outgroup because they belong to the same subfamily of Phocidae (Slade et al. 1994). The southern elephant seal data had been sampled from the following locations: Marion Island (*n* = 50), King George Island (*n* = 23), Macquarie Island (*n* = 53), Elephant Island (*n* = 12), Falkland Islands (*n* = 16), Península Valdés (*n* = 32), Heard Island (*n* = 6), and South Georgia (*n* = 28). We aligned the 12 newly generated sequences and the 201 published sequences using Geneious Prime (Kearse et al. 2012).

### Genetic analyses

Uncorrected pairwise distances were calculated from sampled sequences of the mitochondrial control region using MEGA X (Kumar et al. 2018). We used pairwise deletion to account for gaps in sequences. We used DnaSP version 6.12 (Rozas et al. 2017) to calculate the number of segregating sites, number of haplotypes and haplotype diversity, nucleotide diversity and average nucleotide differences, and neutrality tests using Fu’s *F*_S_ (Fu 1997). Rarefaction was used to correct for unequal sample sizes by comparing haplotype richness between the samples from Davis Base and from all other populations. The sampled haplotype sequences were rarefied to generate the expected haplotype richness using the function *rarefy* (Hurlbert 1971; Heck et al. 1975) in R package vegan (Oksanen et al. 2020).

To visualize the relationships among mitochondrial haplotypes, we constructed a median-joining haplotype network using the software package POPART (Leigh and Bryant 2015). We assigned each seal sequence to one of nine geographical locations: Marion Island (MR), King George Island (KG), Macquarie Island (MQ), Elephant Island (EI), Falkland Islands (FI), Península Valdéz (PV), Heard Island (HD), South Georgia (SG), and Davis Base. The sequence data collected from GenBank were assigned locations according to where they were collected for the referenced study.

### Animal tracking

We immobilized the 12 seals as part of an integrated oceanography and animal behaviour study (McMahon et al. 2021). From each seal, we took morphometric measurements including standard body length, maximum girth, and weight (Field et al. 2002). Each seal was anaesthetised using Zoletil 100, or available combinations of Tiletamine and Zolazepam (McMahon et al. 2000). We then attached identification tags to the hind flippers using Dalton Jumbo Robotags (Wilkinson and Bester 1997). We tracked the 12 seals over a year of their lifecycle from post-moulting to the next moulting haul-out season the following year (Table 1). Conductivity-Temperature-Depth Satellite Data Relay Loggers (CTD-SRDL, Sea Mammal Research Unit, University of St Andrews, UK) were glued to the top of the seal’s head for improved satellite reception and transmission at sea (Horning et al. 2019). The tag attachments to the seals did not affect their reproduction or survival patterns (McMahon et al. 2008). Tracking data received from the tags, such as location and diving behaviour, were transmitted via the ARGOS satellite network (Myers et al. 2006; Henderson et al. 2020).

**Table 1.** Male southern elephant seals (*Mirounga leonina*) with tracking tags. Weights of 54_SES and 64_SES were not taken. Data were collected at Davis Base, Antarctica (68° 34′ 36″ S, 77° 58′ 03″ E).

| Sample Label | Age Class | Mass (kg) | Tag date (d-m-y) |
| --- | --- | --- | --- |
| 42_SES | Subadult | 338 | 17-01-2016 |
| 44_SES | Subadult | 320 | 15-02-2016 |
| 46_SES | Subadult | 406 | 17-02-2016 |
| 48_SES | Subadult | 392 | 20-02-2016 |
| 50_SES | Subadult | 299 | 07-02-2017 |
| 52_SES | Subadult | 309 | 16-02-2017 |
| 54_SES | Juvenile | NA | 16-02-2017 |
| 56_SES | Subadult | 265 | 16-02-2017 |
| 58_SES | Subadult | 345 | 20-02-2017 |
| 60_SES | Subadult | 237 | 21-02-2017 |
| 62_SES | Subadult | 380 | 23-02-2017 |
| 64_SES | Juvenile | NA | 25-02-2017 |

We analysed the tracking data using the R package foieGras (Jonsen and Patterson 2020). We first used the *fit_ssm* function to fit a continuous-time correlated random walk state-space model (SSM; Jonsen et al. 2020) to the ARGOS satellite-derived locations. This model accounted for well-known measurement errors in the ARGOS locations and prredicted locations at regular 12-h time intervals along the seal tracks (as per Jonsen et al. 2019). We then used the *fit_mpm* function to fit a movement persistence model (Jonsen et al. 2019) to the predicted locations to infer changes in the seals’ movement behaviour, possibly arising in response to stimuli such as changes in prey density or ice concentration, along their estimated tracks. Movement persistence (γ_*t*_) is the autocorrelation in both speed and direction (scaled from 0 to 1) between successive displacements along a movement pathway. Low γ_*t*_ values represent low speed and/or directionality that are typical of resident or area-restricted searching behaviours, whereas high γ_*t*_ values represent higher speed and/or directionality that are typical of directed travel associated with dispersal or migration. Using the SSM-predicted locations, we also calculated the following track summary statistics: maximum displacement from deployment location; total deployment duration; maximum displacement scaled by deployment duration; and path tortuosity (mean vector of turning angles along each seal’s track).

## Results

### Population genetic diversity

Nucleotide sequences of the mitochondrial control region (348 bp), sampled from the 12 SES at Davis Base, showed extensive divergence between sample 48_SES and the other 11 samples (the latter were all closely related to each other). Analysis of these 12 sequences and 191 published southern elephant seal sequences revealed a close relationship between sample 48_SES and sequences from Macquarie Island seals. The largest pairwise genetic distance (0.065 substitutions per site) is seen between seals from Macquarie Island and Península Valdés (Table 2). This is expected, given that the two locations are geographically the farthest apart. The pairwise distance between the Davis Base and Macquarie Island seals (0.049 substitutions per site) is greater than that between the Davis Base seals and other population groups. This confirms the divergence between sample 48_SES and the other 11 Davis Base seals.

**Table 2.**
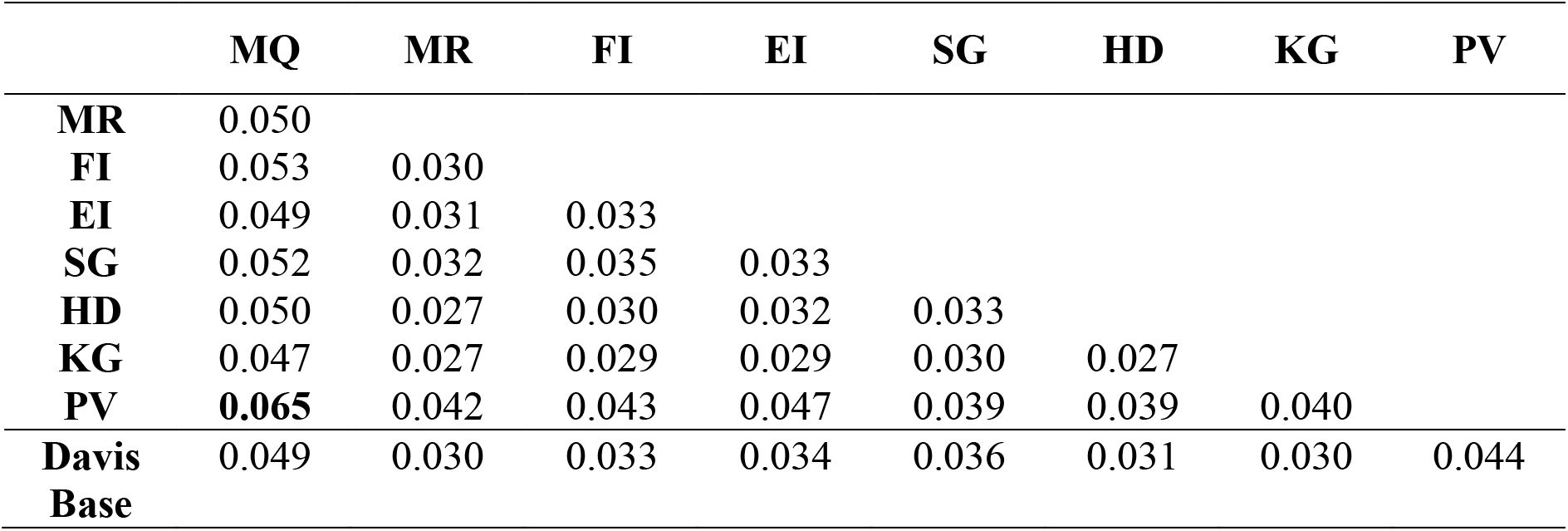
Pairwise mitochondrial genetic distances (nucleotide substitutions per site) between seals from eight population groups and Davis Base. Estimated standard errors for all values were approximately 0.01. Bold font denotes the largest nucleotide pairwise distance, between seals from Macquarie Island and Península Valdés. Location groups are: Macquarie Island (MQ), Marion Island (MR), Falkland Islands (FI), Elephant Island (EI), South Georgia (SG), Heard Island (HD), King George Island (KG), Península Valdés (PV).

We identified a total of 65 haplotypes in the combined data set of 203 mitochondrial sequences (Table 3). From the 12 seals from Davis Base, 11 haplotypes were identified. A single haplotype was carried by 26 individual seals that were previously sampled from King George Island, Marion Island, Falkland Islands, Heard Island, and one individual from Elephant Island. Six haplotypes from Davis Base seals were shared with those from Marion Island, and three haplotypes with those from the Falkland Islands. The final Davis Base haplotype grouped with Macquarie Island, as described above. The seals from Davis Base had high haplotype diversity (0.99 ± 0.04) and had higher nucleotide diversity (2.52 ± 0.004) than the seals from the other populations (Macquarie Island, Marion Island, Heard Island, King George Island, and Península Valdés). Owing to the small number of seal samples from Davis Base, haplotype richness for all population groups was corrected for sample size, and Davis Base seals were elevated compared with all other populations, as well as for the average number of nucleotide differences.

**Table 3.** Statistical summary data for all southern elephant seal populations, including Davis Base seals. Statistics included are the following: the number of individuals (*N*), haplotype diversity (*H*_d_), nucleotide diversity (*π*), average number of nucleotide differences (*k*), Fu’s *F*_S_ statistic (*F*_S_), and haplotype richness based on a sample *n* = 12 (*r*_(12)_).

| Population | $N$ | $H_d \pm SD$ | $\pi (\%) \pm SD$ | $k$ | $F_S$ | $r_{(12)}$ |
| --- | --- | --- | --- | --- | --- | --- |
| Macquarie Island | 53 | $0.92 \pm 0.02$ | $2.50 \pm 0.002$ | 5.73 | -1.00 | 7.24 |
| Marion Island | 50 | $0.98 \pm 0.01$ | $2.51 \pm 0.002$ | 5.71 | -24.87 | 7.53 |
| Falkland Islands | 16 | $1.00 \pm 0.02$ | $3.16 \pm 0.002$ | 7.24 | -10.51 | 10.40 |
| Elephant Island | 12 | $1.00 \pm 0.03$ | $3.29 \pm 0.002$ | 7.53 | -6.13 | 8.00 |
| South Georgia | 28 | $0.99 \pm 0.01$ | $2.84 \pm 0.002$ | 6.51 | -18.42 | 9.13 |
| Heard Island | 6 | $1.00 \pm 0.10$ | $2.42 \pm 0.005$ | 5.53 | -1.96 | 4.00 |
| King George Island | 23 | $0.91 \pm 0.05$ | $1.03 \pm 0.002$ | 2.22 | -9.55 | 8.56 |
| Península Valdés | 32 | $1.00 \pm 0.27$ | $0.58 \pm 0.002$ | 1.33 | -1.22 | 2.00 |
| Davis Base | 12 | $0.99 \pm 0.04$ | $2.52 \pm 0.004$ | 8.73 | -3.31 | 11.00 |
| All combined | 203 | $0.96 \pm 0.01$ | $1.96 \pm 0.001$ | 4.20 | -60.75 | |

### Haplotype network

We assigned traits (locations) to the 12 sequences from Davis Base seals based on their grouping with the known locations of seal sequences from GenBank (Fig. 2). From the combined analysis of all population groups and the Davis Base seals, 12 haplotypes were unique to the Macquarie Island stock. One Davis Base seal (48_SES) shared a haplotype with four seals from Macquarie Island. Eleven haplotypes were unique among seals from Marion Island. Out of the 11 Marion Island haplotypes, four haplotypes were shared by seals from the Falkland Islands, three shared by seals from each of Elephant Island, Heard Island, and King George Island, and two shared by seals from South Georgia. Islands in the South Atlantic Ocean region mostly shared haplotypes with one another, with only 7.69% of haplotypes shared with populations on islands in the South Indian Ocean. We found no common haplotypes between Macquarie Island (South Pacific Ocean), the South Atlantic Ocean, and South Indian Ocean islands.

**Fig. 2.**
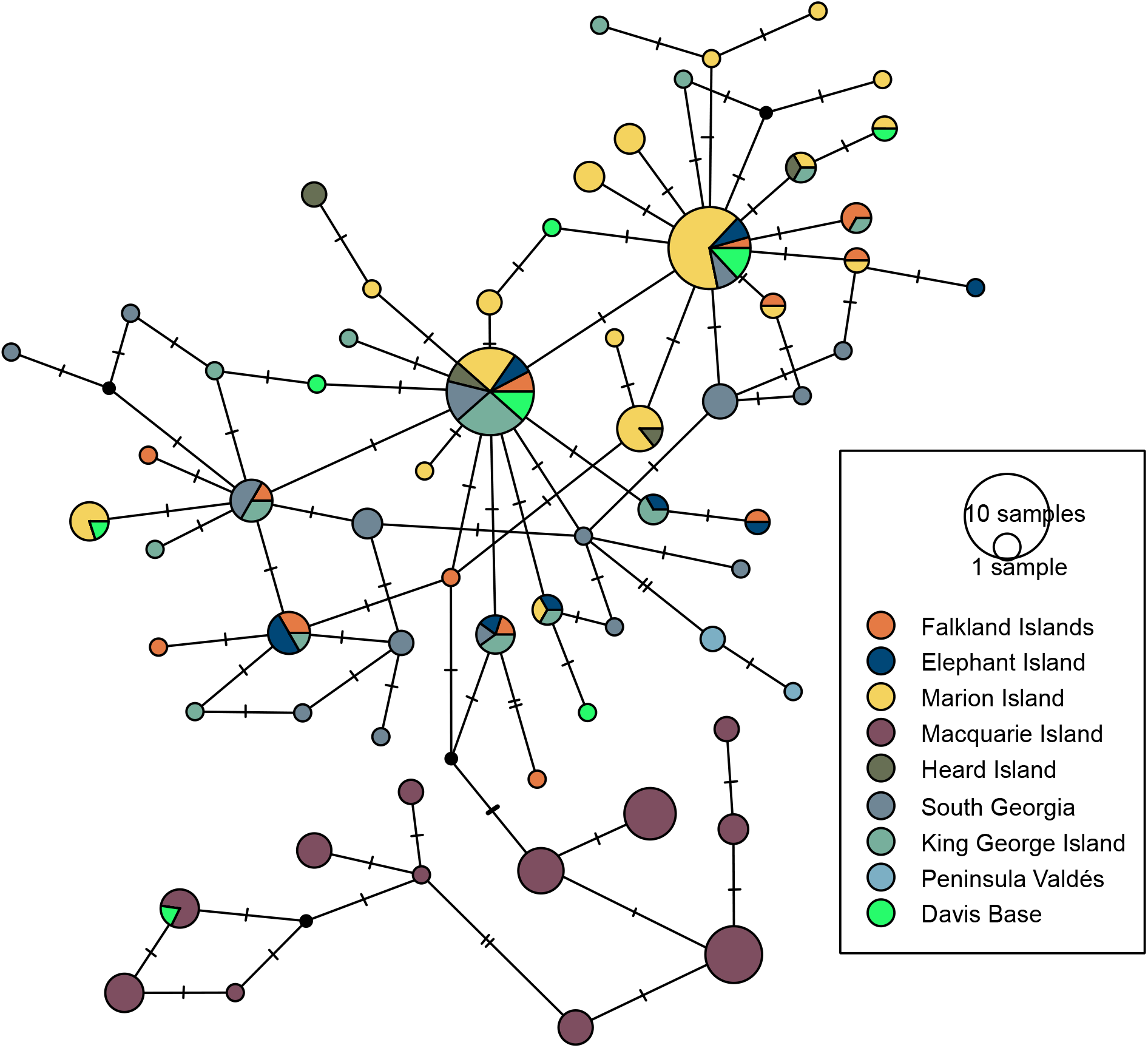
Median-joining network of mitochondrial haplotypes shared between southern elephant seals from Davis Base and all other major populations. Sizes of circles represent the number of individuals per haplotype. Each colour in the pie charts represents a unique population group. Davis Base samples are indicated as light green. The Weddell seal outgroup is shown by a red arrow and is separated by 80 mutations. Black dots represent unobserved haplotypes that have one mutational step from adjacent haplotypes. Hatch marks on lines connecting haplotypes indicate mutations.

### Animal tracking analyses

We analysed track summaries, estimated move persistence, and predicted locations of the 12 seal samples from Davis Base (Table 4). Track summaries indicated that the average maximum displacement of the seals from Davis Base was 1,330 km, with 44_SES travelling the greatest distance (3,056 km). The average maximum displacement scaled by duration was found to be 8.10 km/day, with the greatest being 14.46 km/day (62_SES), and the smallest being 1.30 km/day (seal 60). The average path tortuosity was found to be 0.08, with 58_SES having the highest path tortuosity (0.19), and 44_SES and 54_SES with the lowest (0.02).

**Table 4.** Southern elephant seal track summaries by maximum displacement from the track start (*D*_max_), tracking deployment days (duration), maximum displacement scaled by deployment duration (*D*_max_.*d*_t_), and path tortuosity 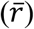.

| Sample | $D_{\max}$ (km) | duration (days) | $D_{\max}.d_t$ (km/day) | $\bar{r}$ |
| --- | --- | --- | --- | --- |
| 42_SES | 634.57 | 143.0 | 4.44 | 0.07 |
| 44_SES | 3055.92 | 270.0 | 11.32 | 0.02 |
| 46_SES | 1013.30 | 74.0 | 13.69 | 0.06 |
| 48_SES | 2967.28 | 305.0 | 9.73 | 0.04 |
| 50_SES | 2584.35 | 246.0 | 10.51 | 0.13 |
| 52_SES | 691.64 | 59.5 | 11.62 | 0.10 |
| 54_SES | 250.99 | 57.5 | 4.37 | 0.02 |
| 56_SES | 2234.81 | 216.0 | 10.35 | 0.05 |
| 58_SES | 142.90 | 48.0 | 2.98 | 0.19 |
| 60_SES | 28.66 | 22.0 | 1.30 | 0.14 |
| 62_SES | 2205.88 | 152.5 | 14.46 | 0.06 |
| 64_SES | 147.04 | 60.5 | 2.43 | 0.05 |
| **Average** | 1329.78 | 137.8 | 8.10 | 0.08 |

The sampled seals showed high move persistence during their outbound trips into open waters, and low move persistence when approaching areas with high sea-ice coverage or when returning to breeding and moulting areas (Fig. 3). Predicted locations have indicated visits to Kerguelen Islands by 44_SES, 50_SES, 56_SES, and 62_SES (Fig. 4). Seal 48_SES made visits to Crozet Islands and appeared to spend the majority of its journey around those islands. The predicted locations also indicated that five individuals did not venture out of the Davis Base ice shelf regions. This might be due to seal mortality or detachment of the tags.

**Fig. 3.**
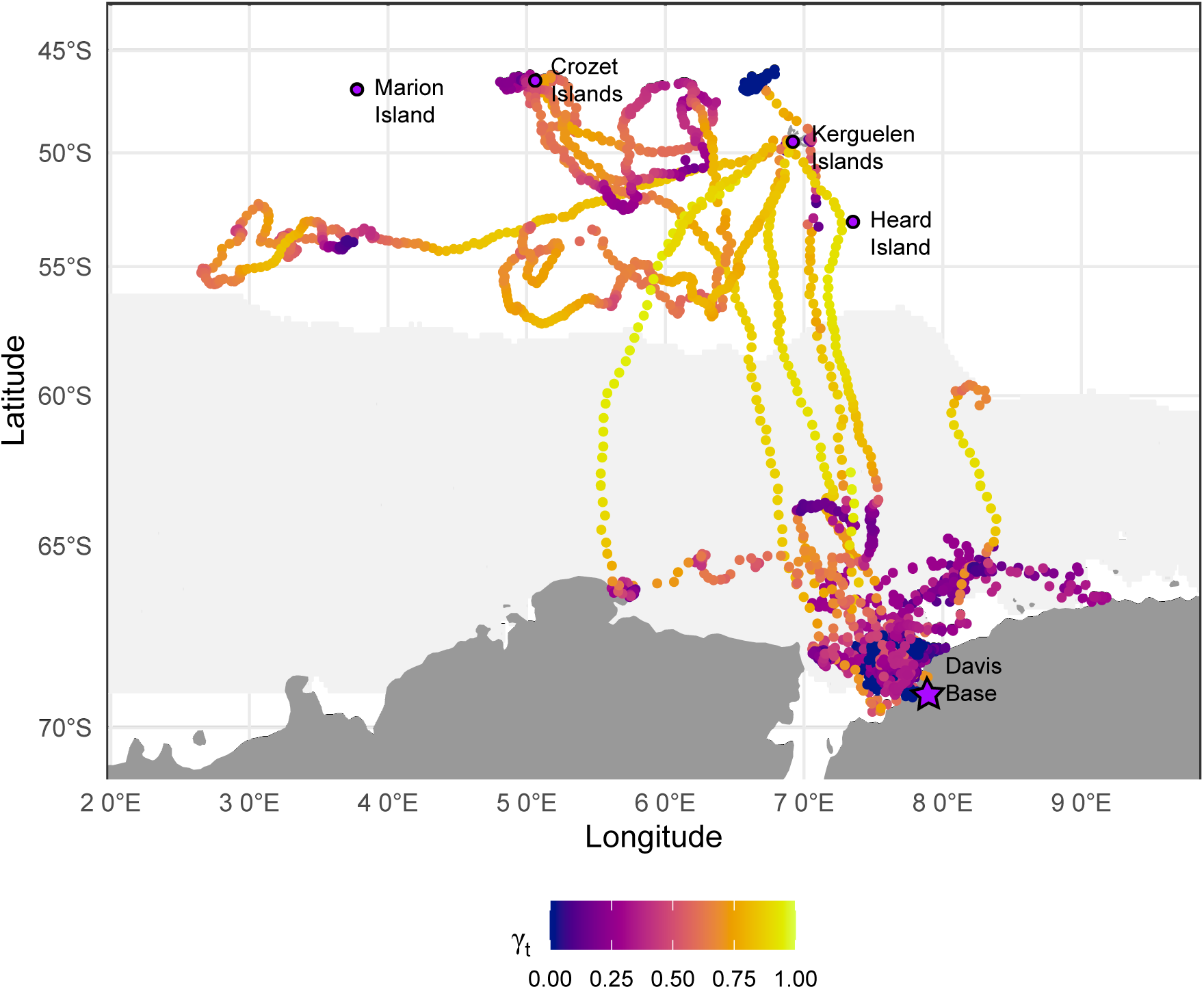
Map of sampled seals’ move persistence by measure of path tortuosity. γ_t_ values approaching 1.00 (yellow) indicate relatively fast, directed movement. γ_t_ values approaching 0.00 (navy) indicate high tortuosity and slow movement. Davis Base is indicated by the purple star. Main breeding islands are represented by purple circles. Ice concentrations (maximum extent of sea ice coverage >15% during winter) are represented by light grey along the coastline.

**Fig. 4.**
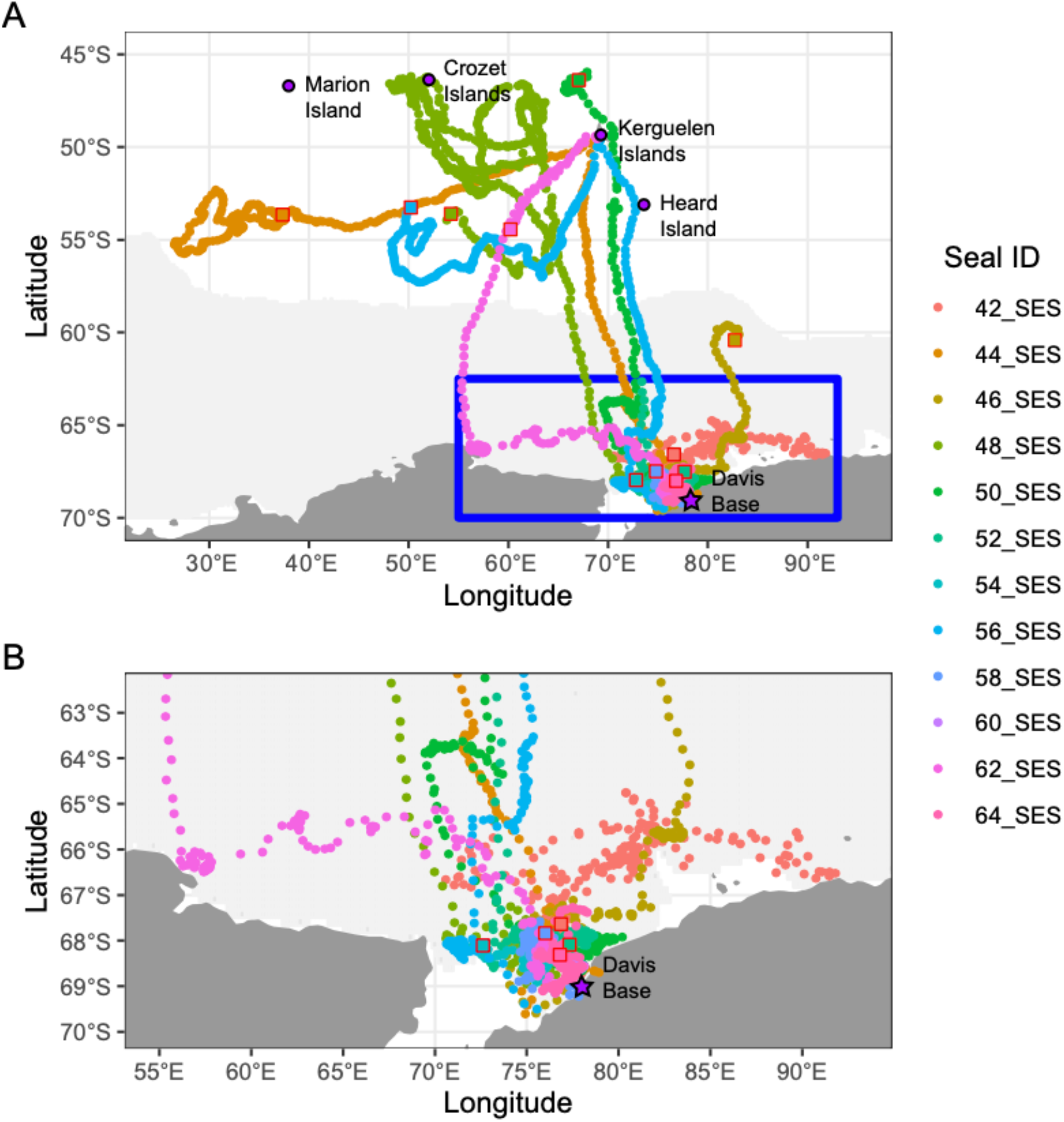
(A) State-space model predicted locations of all sampled seals. Seal 60_SES failed to converge on predicted locations due to small scales and low contrasts of movement. The blue box indicates a narrowed view of the cluster of seal tracks close to Davis Base as seen in (B). The end track location of each seal is shown by a red outlined square. Davis Base is indicated by the purple star. Breeding islands are represented by purple circles. Ice concentrations (maximum extent of sea ice coverage >15% during winter) are represented by light grey along the coastline.

## Discussion

By combining genetic and animal tracking data, we determined the natal locations and at-sea movements of moulting seals at Davis Base, Antarctica. Our analysis of genetic data revealed a mixed sample of likely natal locations from the 12 Davis Base seals, including South Georgia, Macquarie Island, Marion Island, King George Island, and the Falkland Islands. We also identified two distinct lineages, representing three of the four main breeding stocks (Macquarie Island, South Atlantic Ocean islands, and South Indian Ocean islands). Our tracking data showed that all seals, despite their varied lineages, remained within the vicinity of Crozet Island, Kerguelen Islands, and Heard Island for the duration of the tracking study. Seals that showed short displacements and less move persistence travelled more frequently around areas of high ice concentration. Overall, our integrated genetic and telemetry analyses provided longer-term insights into the seals’ natal affiliations and migration strategies, and additional short-term information on their annual life-cycle movements and foraging ecology.

The genetic data from 12 SES collected at Davis Base suggested high nucleotide diversity and haplotypes shared with elephant seals from other breeding colonies within the South Atlantic Ocean, South Indian Ocean, and South Pacific Ocean. However, given the small sample size, our study might not be representative of the larger population of all male SES at Davis Base. Replication of this study with a larger sample size would assist in describing more comprehensively the genetic make-up of this population. Our key findings are that 11 of the 12 seals had unique mitochondrial DNA haplotypes and one seal shared a haplotype with seals from Macquarie Island. Migration and mixing between the Kerguelen and Macquarie stocks has not been observed previously. Seals from Macquarie Island form a monophyletic group and are one of the four major breeding stocks of SES. The other 11 seals from Davis Base showed mixed affinities to island populations of two other major breeding stocks: those of the South Indian Ocean (Marion Island and Heard Island), and South Atlantic Oceans (King George Islands, Falkland Islands, and Elephant Island). These two breeding stocks display extensive gene flow and intermixing, as shown by the sharing of haplotypes between stocks. Two of these individuals (56_SES and 64_SES) shared a known haplotype with seals from Marion Island only.

### Animal tracking analyses

The male seals from our study spent more time in areas of high ice concentration and typically involved intense search behaviours exemplified by highly tortuous movements. Similar patterns have been observed previously in male seals that adopted sea-ice foraging strategies rather than pelagic foraging (Labrousse et al. 2017; Rodríguez et al. 2017; Jonsen et al. 2019; Hindell et al. 2021). Moreover, the preferences of foraging locations along the Kerguelen Plateau were shown by seal 50_SES, which had less move persistence near Kerguelen Islands. Five other seal tracks (44_SES, 48_SES, 50_SES, 56_SES, and 62_SES) showed patterns of inter-island movements by outbound trips made towards the Kerguelen Plateau and Crozet Islands, which might indicate pelagic foraging trips away from the ice shelf areas. Similarly, seal 48_SES showed less move persistence and had the second-longest maximum displacement per day around Crozet Islands (which is approximately 6,900 km from its natal affinity, Macquarie Island). This suggests a long-distance dispersal event to gain foraging (and potentially future breeding) advantage away from the seal’s natal location.

SES are philopatric to breeding and foraging sites, so long-distance dispersal events are rare, particularly for a juvenile seal (Hindell and McMahon 2000; Reisinger and Bester 2010). Our findings show that the only two male juvenile seals, out of the 12 seals included in this study, remained close to the ice shelf region to forage close to Davis Base, which was indicated by the reduced move persistence. Seals that remain close to the ice shelf region might have easier access to sea-ice polynyas where their prey (myctophids and ice fish) may be more easily found (Labrousse et al. 2017). However, a previous study found that a juvenile female from Macquarie Island was sighted at Peter 1 Øy, which is approximately 5,200 km to the east (Hindell and McMahon 2000). This finding corresponds with that observed here of 48_SES (Macquarie Island natal affinity), which foraged around Crozet Islands, as discussed above. The parallel finding between this study and previous studies might indicate a lack of resource availability around Macquarie Island where the population has been in decline in recent decades (McMahon et al. 2005a), which could be forcing seals from this location to disperse farther to find food.

### Changes to foraging site and the consequences on populations

The Kerguelen Plateau is a popular foraging location for male SES. However, there are high levels of predation on SES here (Hindell et al. 2021), which might have caused terminations of seal tracking signals. The Kerguelen Plateau lies within the Antarctic Polar Front boundary, which is an area of elevated productivity for most Antarctic marine species due to the decrease in water temperature down to 2°C at 200 m, and the distribution of water masses and associated abundance of primary producers (O’Toole et al. 2014; Cristofari et al. 2018). SES will often forage on the Antarctic Continental Shelf and Polar Front, and because they are a deep-diving species, sea-surface temperatures have less of an effect on their diving behaviours (Hindell et al. 1991; O’Toole et al. 2014). Alternatively, previous studies have found that seals have greater foraging opportunities in colder waters, particularly along the Antarctic shelf; temperatures slow down the movements of prey, allowing their capture with less energy expenditure from the seals (Bailleul et al. 2007).

Popular foraging locations around the Kerguelen Plateau, such as Crozet Islands and Kerguelen Islands, have been reported to show either a slight increase or a stabilization in SES population numbers over the last decade (Guinet et al. 2004). The stabilized population numbers might have been caused by the seals’ inter-island movements and foraging between islands in the Kerguelen province, including Marion Island, Crozet Islands, Kerguelen Islands, and Heard Island (Oosthuizen et al. 2011). However, despite showing philopatry to foraging sites, the seals’ foraging strategies and movements might shift based on the changes to oceanic conditions that will influence where resources are available (Bailleul et al. 2007). Moreover, poor foraging success by females has led to a decrease in first-year pup survival and would thus reduce reproductive success in populations such as Macquarie Island (Arnbom et al. 1997; McMahon et al. 2003; Clausius et al. 2017; Mestre et al. 2020). As oceanic conditions continue to change with climate, the entire marine ecosystem will also shift. Therefore, the lack of resources around Macquarie Island is likely to continue causing decreases in population size (McMahon et al. 2005a; Clausius et al. 2017).

### Population declines impacted by climate change

Broad climate events such as El Niño can affect the oceanic structure of the Southern Ocean. The Southern Ocean ecosystem relies heavily on phytoplankton and krill abundance for other species’ survival, previously reported for Antarctic whales, king penguins, and SES (McIntyre et al. 2014; Cristofari et al. 2018; Bestley et al. 2020; Rogers et al. 2020; Agrelo et al. 2021; Volzke et al. 2021). Over the last four decades, the global southern elephant seal population has seen dramatic declines in some populations due to the changes in food availability (McMahon et al. 2005a; Volzke et al. 2021). Populations of the four main breeding stocks, South Georgia, Kerguelen Islands and Heard Island, Macquarie Island, and Peninsula Valdés, have all decreased since the 1970s. However, presently all the populations bar the Macquarie population are either stable or increasing (McMahon et al. 2005b; Hindell et al. 2016). A more thorough understanding of seal foraging behaviour, the selection and variations in foraging sites, and how this is expressed and transmitted within a population is central to understanding how foraging site selection affects population growth and ultimately population variability.

In summary, we were able to identify the natal locations of the 12 Davis Base seals through genetic data. We combined findings from genetic data with satellite telemetry tracking data and identified that the majority of the seals are spending most of the time foraging along the Kerguelen Plateau. Our data suggest that a seal from Macquarie Island has travelled a long distance, probably to gain a foraging advantage over its conspecifics. Long-range migrations and dispersal to distant feeding grounds might be one way for seals to maximize foraging efficiency, which may affect population growth rates through changes in survival and reproduction.

## Acknowledgements

We thank the Integrated Marine Observing System (IMOS) team for collecting and providing the elephant seal samples and tracking data. IMOS is a national collaborative research infrastructure, supported by the Australian Government and operated by a consortium of institutions as an unincorporated joint venture, with the University of Tasmania as Lead Agent. S.Y.W.H. was funded by the Australian Research Council. Colleagues in the Molecular Ecology, Evolution, and Phylogenetics Lab provided advice and support throughout this project.

## Author contributions

C.R.M. conducted Antarctic fieldwork and collected data. M.C. did the molecular lab work, conducted molecular and tracking data analyses, and wrote the first draft of the manuscript. I.J. provided guidance on and contributed to tracking data analysis. S.Y.W.H, C.R.M., and M.d.B supervised the research project. All authors revised and contributed to the writing of the final manuscript.

